# A theory of working memory without consciousness or sustained activity

**DOI:** 10.1101/093815

**Authors:** Darinka Trübutschek, Sébastien Marti, Andrés Ojeda, Jean-Rémi King, Yuanyuan Mi, Misha Tsodyks, Stanislas Dehaene

## Abstract

Working memory and conscious perception are thought to share similar brain mechanisms, yet recent reports of non-conscious working memory challenge this view. Combining visual masking with magnetoencephalography, we demonstrate the reality of non-conscious working memory and dissect its neural mechanisms. In a spatial delayed-response task, participants reported the location of a subjectively unseen target above chance-level after a long delay. Conscious perception and conscious working memory were characterized by similar signatures: a sustained desynchronization in the alpha/beta band over frontal cortex, and a decodable representation of target location in posterior sensors. During non-conscious working memory, such activity vanished. Our findings contradict models that identify working memory with sustained neural firing, but are compatible with recent proposals of ‘activity-silent’ working memory. We present a theoretical framework and simulations showing how slowly decaying synaptic changes allow cell assemblies to go dormant during the delay, yet be retrieved above chance-level after several seconds.

## Introduction

Prominent theories of working memory require information to be consciously maintained (Baars and Franklin, 2003; Baddeley, 2003; Oberauer, 2002). Conversely, influential models of visual awareness hold information maintenance as a key property of conscious perception, highlighting synchronous thalamocortical activity (Tononi and Koch, 2008), cortical recurrence (Lamme and Roelfsema, 2000), or the sustained recruitment of parietal and dorsolateral prefrontal regions (i.e., the same areas as in working memory; Naghavi and Nyberg, 2005) in a global neuronal workspace (Dehaene and Changeux, 2011; Dehaene and Naccache, 2001). Experimentally, non-conscious priming only lasts a few hundred milliseconds (Dupoux et al., 2008; Greenwald et al., 1996) and unseen stimuli typically fail to induce late and sustained cerebral responses (Dehaene et al., 2014). Conscious perception, in contrast, exerts a durable influence on behavior, accompanied by sustained neural activity (King et al., 2014; Salti et al., 2015; Schurger et al., 2015). The hypothesis of an intimate coupling between conscious perception and working memory is thus grounded in theory and supported by numerous empirical findings.

Recent behavioral and neuroimaging evidence, however, has questioned this prevailing view by suggesting that working memory may also operate non-consciously. Unseen stimuli may influence behavior for several seconds (Bergström and Eriksson, 2015; Soto and Silvanto, 2014). Soto and colleagues (Soto et al., 2011), for instance, showed that participants recalled the orientation of a subjectively unseen Gabor cue above chance-level after a 5s-delay. Functional magnetic resonance imaging suggests that prefrontal activity may underlie such non-conscious working memory (Bergström and Eriksson, 2014; Dutta et al., 2014).

The verdict for non-conscious working memory is far from definitive, however. Delayed performance with subjectively unseen stimuli was barely above chance (Soto et al., 2011) and could have arisen from a small percentage of errors in visibility reports, with subjects miscategorizing a seen target as unseen (miscategorization hypothesis). Alternatively, participants could have ventured a guess about the target as soon as it appeared and consciously maintained this early guess (conscious maintenance hypothesis). Many priming studies have shown that fast guessing results in above-chance objective performance with subjectively unseen stimuli (Merikle, 2001). The observed blindsight effect would then reflect a normal form of conscious working memory (Stein et al., 2016). Lastly, even if non-conscious working memory had been convincingly demonstrated, no mechanistic account has been offered for it. Here, we set out to address these issues by testing the reality of non-conscious working memory, interrogating the link between conscious perception and working memory, and identifying the brain mechanisms associated with conscious and non-conscious working memory maintenance.

## Results

We combined magnetoencephalography (MEG) with a spatial masking paradigm to assess working memory performance under varying levels of subjective visibility (Figure 1A and Methods). On 80% of the trials, a target square was flashed in 1 of 20 locations and then masked. Subjects were asked to localize the target after a variable delay (2.5 – 4.0s) and to rate its visibility on a scale from *1* (not seen) to *4* (clearly seen). On the remaining 20% of trials, the target was omitted, allowing us to contrast brain activity between target-present and -absent trials. A visible distractor square was presented 1.5s into the delay period on half the trials, challenging participants’ resistance to distraction. In addition to this working memory task, subjects also completed a perception-only control condition without the delay and target-localization periods (perception task), enabling us to isolate brain activity specific to conscious perception and to investigate its link with working memory.

**Figure 1.**
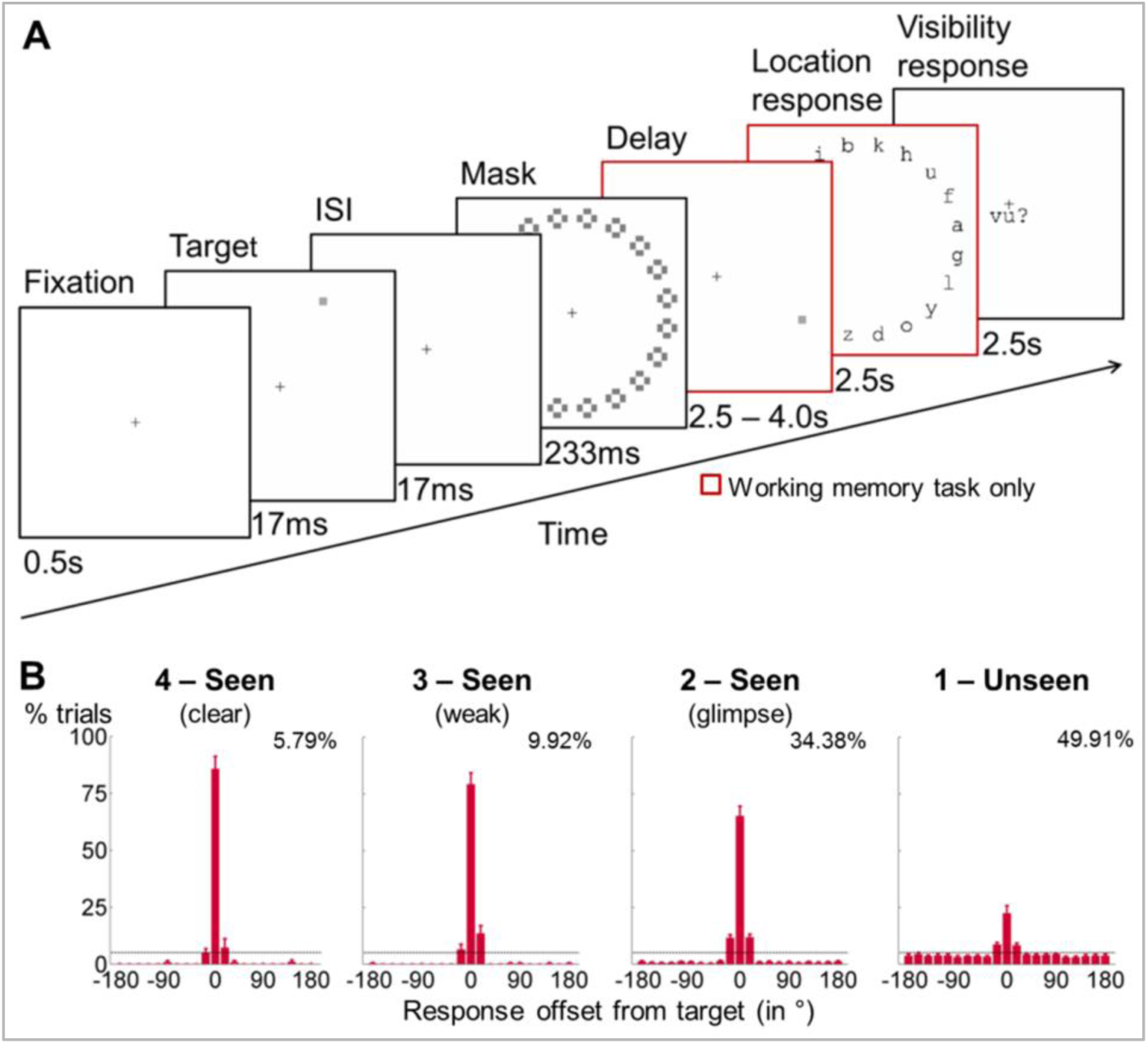
**General experimental design and behavioral performance in the working memory task**. **(A)** Experimental design. A subsequently masked target square was flashed in 1 out of 20 positions. Subjects were asked to report this location after a delay of up to 4s and to rate the visibility of the target on a 4-point scale. A visible distractor square with features otherwise identical to the target was shown on 50% of the trials during the retention period (at 1.75s). In a perception-only control condition, the maintenance phase and location response were omitted, and subjects assessed the visibility of the target immediately after the mask.**(B)** Spatial distributions of forced-choice localization performance in the working memory task (experiment 1; 0 = correct target location; positive = clockwise offset). Error bars indicate standard error of the mean (SEM) across subjects. The horizontal, dotted line illustrates chance-level at 5%. Percentages show proportion of targetpresent trials from a given visibility category. Due to low number of trials in individual visibility ratings *2, 3*, and 4, all *seen* categories were collapsed for analyses.

### Behavioral maintenance and shielding against distraction

As subjective visibility of the target increased from glimpsed (visibility = *2*) to clearly seen (visibility = *4*), participants became more accurate in identifying the exact target position (Figure 1B, all pair-wise comparisons: *p* < .05). Overall, subjects reported the original target location with high accuracy on seen trials (collapsed across visibility ratings *> 1*; *M* = 69.1%, *SD* = 17.4%; chance = 5%; *t*(16) = 15.2, *p* < .001) and above chance on unseen trials (rating = 1; *M* = 22.4%, *SD* = 13.8%; *t*(16) = 5.2, *p* < .001). This blindsight remained substantial even after a 4s-delay (*M* = 21.1%, *SD* = 14.7%; *t*(16) = 4.5, *p* < .001).

Spatial distributions of participants’ responses formed a bump around the target (Figure 2A). To correct for small errors in localization, we computed the rate of correct responding (CR) with a tolerance of two positions (+/− 36°) surrounding the target location. In subjects displaying above-chance blindsight (chance = 25%; *p* < .05 in a *χ*^2^-test; *n* = 13), we estimated the precision of working memory as the standard deviation of the distribution within this tolerance interval (Methods). Performance was better on seen than on unseen trials, both in terms of CR (*F*(1, 16) = 198.5, *p* < .001) and precision (*F*(1, 12) = 36.7, *p* < .001). There was neither an effect of the distractor on these measures (all *ps* > .079), nor any significant interactions between distractor and visibility (all *ps* > .251), indicating that distractor presence did not affect retention for seen or unseen targets. Restricting the analysis to trials within one position of the actual target location (+/− 18°) or to the subgroup of 13 subjects included in the MEG analyses did not change these findings qualitatively. These results confirm the observations of previous studies (Soto et al., 2011) with much higher non-conscious performance: Non-conscious information may be maintained for up to 4 seconds and successfully shielded against distraction from a salient visual stimulus.

**Figure 2.**
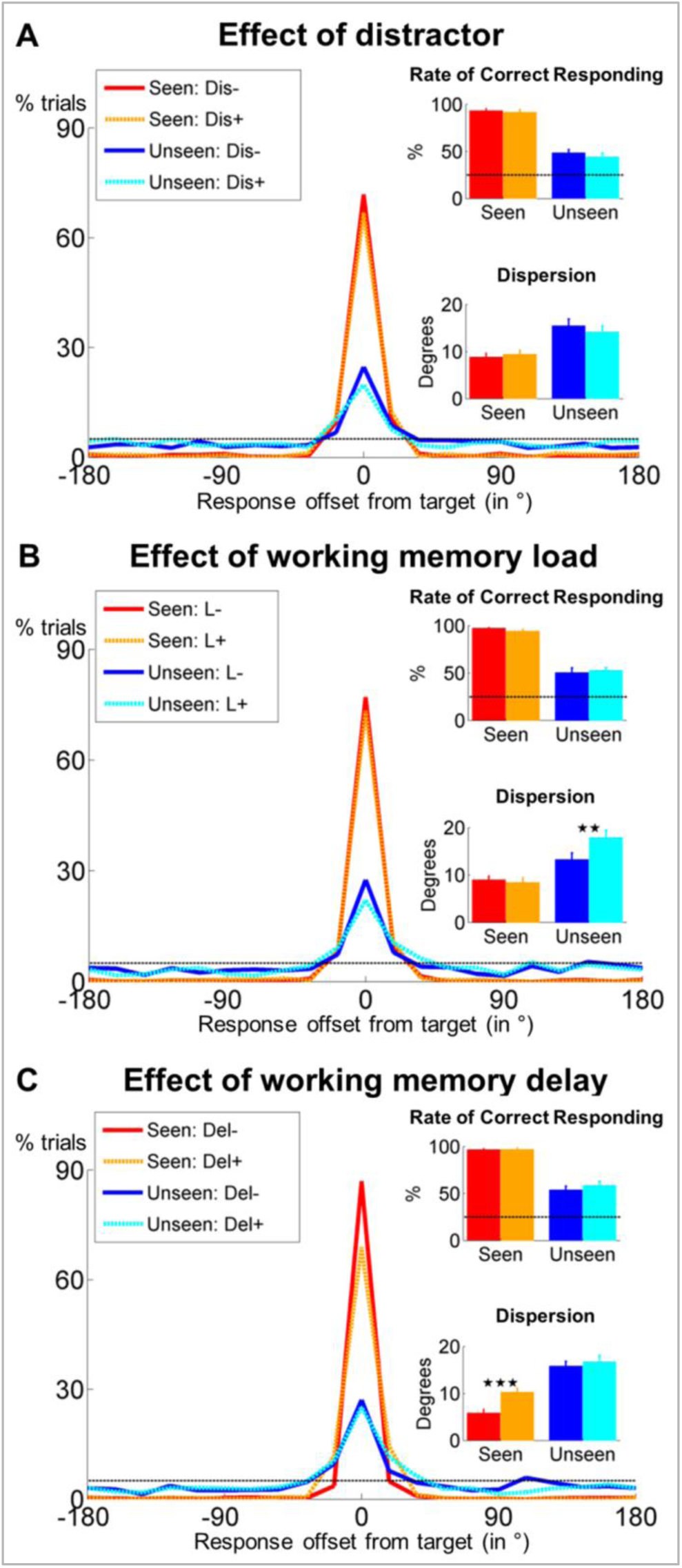
**Behavioral evidence for non-conscious working memory**. Spatial distributions of responses (0 = correct target location; positive = clockwise offset) as a function of visibility and distractor presence **(A)**, conscious working memory load **(B)** and delay duration **(C)**. Insets show rate of correct responding (within +/− 2 positions of actual location) and precision of working memory representation separately for seen and unseen trials. Error bars represent standard error of the mean (SEM) across subjects and horizontal, dotted line indicates chance-level (5%). \**p* < .05, \*\**p* < .01, and \*\*\**p* < .001 in a paired sample *t*-test. Dis = distractor, L = load, Del = delay.

### Resistance to conscious working memory load and delay duration

To probe the similarity between conscious and non-conscious working memory, in a second behavioral experiment with 21 subjects, we examined whether imposing a load on conscious working memory (remembering digits) affected non-conscious performance. On each trial, 1 (low load) or 5 (high load) digits were simultaneously shown for 1.5s, followed by a 1s-fixation period and the same sequence of events (target and mask) as in experiment 1. After a variable delay (0 or 4s), participants had to (1) localize the target, (2) recall the digits in the correct order, and (3) rate target visibility.

Subjects again chose the exact target position with high accuracy on seen trials (*M*= 77.8%, *SD* = 13.9%) and remained above chance on unseen trials (*M* = 25.6%, *SD* = 11.7%; chance = 5%; *t*(18) = 7.6, *p* < .001). As expected, they were better at recalling 1 rather than 5 digits in the correct order (*M* = 93.3% vs. 89.5%, *F*(1, 17) = 4.7, *p* = .045), irrespective of target visibility or delay duration (all *ps* > .135).

Analyzing only the trials with correctly recalled digits, we observed an impact of load on the precision with which target location was retained (*F*(1, 13) = 7.3, *p* = .018). Crucially, load modulated the relationship between precision and visibility (interaction *F*(1, 13) = 8.7, *p* = .011), with no effect on seen (*t*(13) = 0.6, *p* = .561) and a strong reduction of precision on unseen trials (*t*(13) = −3.6, *p* = .004). There was no effect of working memory load on CR (all *ps* > .229; Figure 2B).

Delay duration (0 or 4s) did not influence CR (all *ps* > .082; Figure 2C). It did, however, affect overall precision (*F*(1, 15) = 9.3, *p* = .008; partial η^2^ = .383) and the relationship between precision and visibility (interaction *F*(1, 15) = 5.2, *p* = .037). This interaction was driven by higher precision on no-delay than on 4s-delay trials, exclusively when subjects had seen the target (*t*(15) = −5.7, *p* < .001; unseen trials: *t*(15) = −0.6, *p* = .559).

Overall, these results highlight the replicability and robustness of non-conscious working memory: Even in the presence of a concurrent conscious working memory load, unseen stimuli could be maintained, with no detectable decay as a function of delay. However, the systems involved in the short-term maintenance of conscious and non-conscious stimuli interacted, because a conscious verbal working memory load diminished the precision with which non-conscious spatial information was maintained.

### Similarity of conscious perception and conscious working memory

To probe the mechanisms underlying conscious perception, we turned to the MEG data. The subtraction of the event-related fields (ERFs) evoked by unseen trials from those evoked by seen trials revealed similar topographies for the perception and working memory task (Figure 3A): Starting at ∼300ms and extending until ∼500ms after target onset, a response emerged over right parieto-temporal magnetometers. This divergence resulted primarily from a sudden increase in activity on seen trials (“ignition”) in the perception (*p*_FDR_ < .05 from 384 – 416ms and from 504 – 516ms) and working memory task (*p*_FDR_ < .05 from 328 – 364ms and from 396 – 404ms; Figure 3B). The observed topographies and time courses fall within the time window of typical neural markers of conscious perception, including the P3b (e.g., Del Cul et al., 2007; Salti et al., 2015; Sergent et al., 2005). Consciously perceiving the target stimulus therefore involved comparable neural mechanisms, irrespective of task.

**Figure 3.**
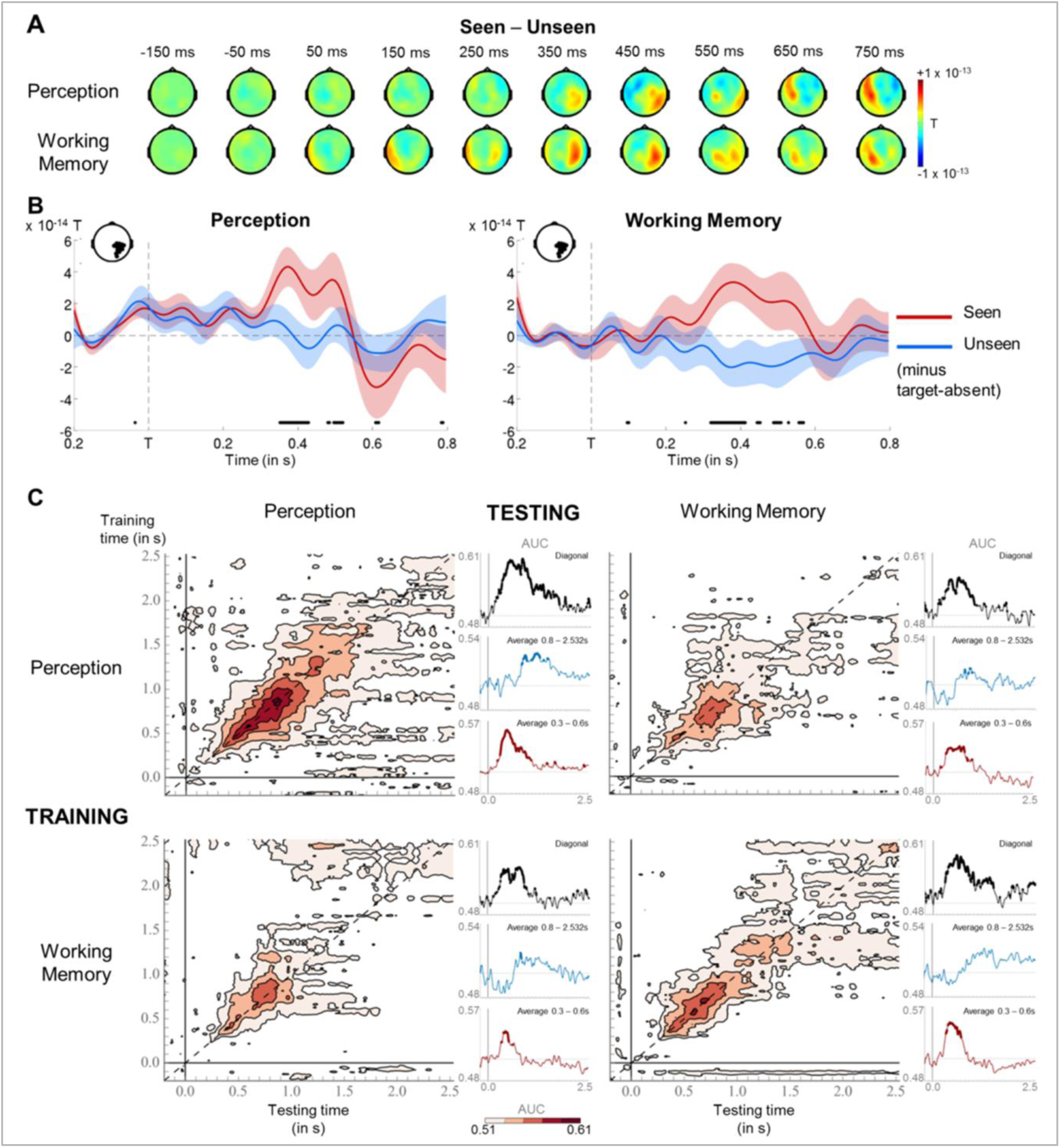
**Neural signatures for conscious perception and maintenance in working memory**. **(A)** Sequence of brain activations (0 – 800ms) evoked by consciously perceiving the target in the perception (top) and working memory (bottom) task. Each topography depicts the difference in amplitude between seen and unseen trials over a 100ms time window centered on the time points shown (magnetometers only). **(B)** Average time courses of seen and unseen trials (0 – 800ms) after subtraction of target-absent trials in a group of parietal magnetometers in the perception (left) and working memory (right) task. Shaded area illustrates standard error of the mean (SEM) across subjects. Significant differences between conditions are depicted with a horizontal, black line (one-tailed Wilcoxon signed-rank test across subjects, uncorrected). For display purposes, data were lowpass-filtered at 8Hz. T = target onset. **(C)** Temporal generalization matrices for decoding of visibility category as a function of training and testing task. In each panel, a classifier was trained at every time sample (y-axis) and tested on all other time points (x-axis). The diagonal gray line demarks classifiers trained and tested on the same time sample. Time courses of diagonal decoding and of classifiers averaged over the working memory maintenance period (0.8 – 2.5s) and over the P3b time window (0.3 – 0.6s) are shown as black, blue, and red insets. Thick lines indicate significant, above-chance decoding of visibility (Wilcoxon signed-rank test across subjects, uncorrected, two-tailed except for diagonal). For display purposes, data were smoothed using a moving average with a window of eight samples. AUC = area under the curve.

We next probed the relationship between conscious perception and information maintenance in working memory. Does the latter reflect a prolonged conscious episode, or does it involve a distinct set of processes recruited only during the retention phase? Multivariate pattern classifiers were trained to predict visibility (seen or unseen) from MEG signals separately for each task. Classification performance was assessed during an early time period (100 – 300ms), the critical P3b time window (300 – 600ms), and the delay period before (0.6 – 1.75s) and after (1.75 – 2.5s) the distractor.

Decoding was comparable in the two tasks (Figure 3C and Table 1): Classification rose sharply between 100 and 300ms and peaked during the P3b time window (all *ps* < .007, except 100 – 300ms in the working memory task, where *p* = .066). It then decayed slowly from ∼1s onward in both tasks, yet remained above chance during the 0.6 – 1.75s interval (all *ps* < .004). Similar time courses were also observed when training in one task and testing for generalization to the other. Though rapidly dropping to chance-level after ∼1s, classifiers trained in the perception task performed above chance during the first three time windows on working memory trials (and vice versa; all *ps* < .020), indicating that, early on, both tasks recruited similar brain mechanisms.

**Table 1.**
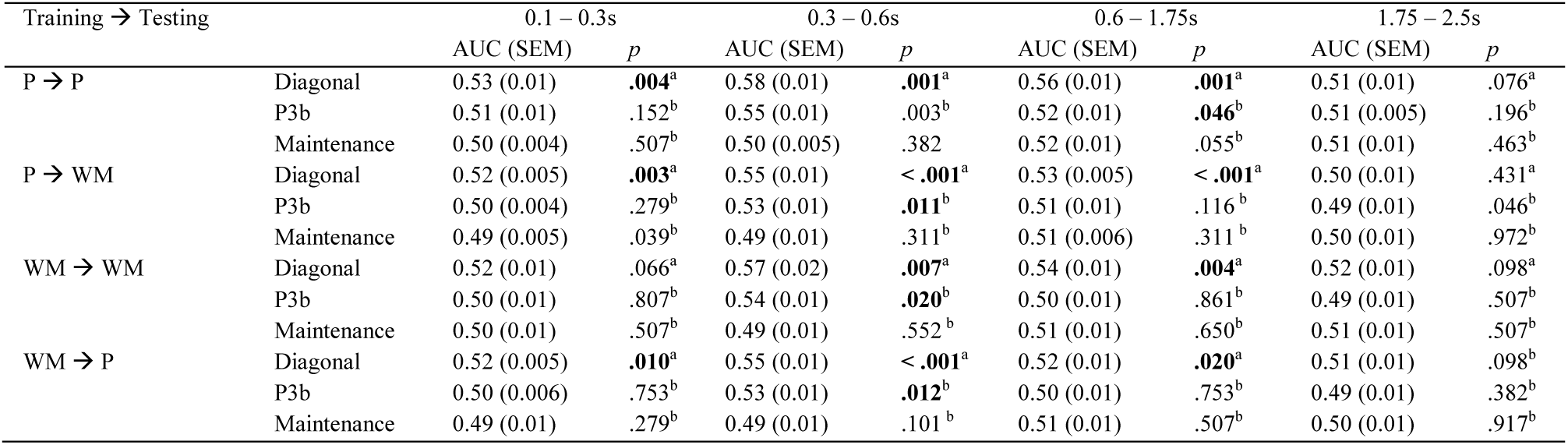
**Statistics for decoding analyses**. Statistics are shown for decoding of visibility category (seen vs. unseen) as a function of task and testing time bin. The first column identifies the respective training and testing sets (P = perception task; WM = working memory task), the second column the training classifiers (Diagonal = diagonal, P3b = 300 – 600ms, Maintenance = 0.8 – 2.5s), that were averaged. Bold numbers indicate above-chance decoding performance (^a^one-tailed, ^b^two-tailed Wilcoxon signed-rank test across subjects). AUC = area under the curve; SEM = standard error of the mean (across participants).

Temporal generalization matrices (King and Dehaene, 2014) were used to evaluate the onset and duration of patterns of brain activity. If working memory were just a prolonged conscious episode, classifiers trained at time points relevant to conscious perception (e.g., P3b) should generalize extensively, potentially spanning the entire delay. Our findings supported this hypothesis only in part. The temporal generalization matrix for the working memory task presented as a thick diagonal, suggesting that brain activity was mainly characterized by changing, but long-lasting patterns. Though failing to achieve statistical significance over the entire 0.6 – 1.75s interval (all *ps* > .116), at a more lenient, uncorrected threshold, classifiers trained during the P3b time window (300 – 600ms) in the working memory task remained weakly efficient until ∼704ms (AUC = 0.53 +/− 0.02, *p*_uncorrected_ =.046). Similarly, classifiers trained during the same time period in the perception task and tested on the working memory task persisted up to ∼988ms (AUC = 0.52 +/− 0.01, *p*_uncorrected_ = .039). Brain processes deployed for the conscious representation of the target were thus partially sustained during the working memory delay. The reverse analysis, in which we trained classifiers during the retention period in the working memory task (0.8 – 2.5s), did not reveal any generalization to the P3b time window in the perception task *(p* > .101).

These results confirm that seeing the target entailed a similar unfolding of neural events in two task contexts: Conscious perception primarily consisted of a dynamic series of partially overlapping information-processing stages, each characterized by temporary, metastable patterns of neural activity. The same neural codes appear to be recruited at the beginning of the maintenance period (up to ∼1s). As such, these findings corroborate previous accounts linking conscious perception to an “ignition” of brain activity (Del Cul et al., 2007; Gaillard et al., 2009; Salti et al., 2015; Sergent et al., 2005) and suggest that, in part, working memory implies the prolongation of a conscious episode, and, in part, a succession of additional processing steps.

### A sustained decrease in alpha/beta power distinguishes conscious working memory

Our focus so far has been on evoked brain activity. However, other reliable neural signatures of conscious perception have been identified in the frequency domain (Gaillard et al., 2009; Gross et al., 2007; Wyart and Tallon-Baudry, 2009). We thus turned to time-frequency analyses and first contrasted seen and unseen trials in both tasks (Figure 4A). Cluster-based permutation analyses singled out a desynchronization in the alpha band (8 – 12Hz) as the principal correlate of conscious perception in the perception task (*p*_clust_ = .009), with seen trials displaying a strong decrease in power compared to the baseline period. Initially left-lateralized in centro-temporal sensors, this effect moved to fronto-central channels and extended between ∼300 and 1700ms. A similar, albeit later (900 – 1700ms) and more bilateral fronto-central, desynchronization was also observed in the beta band (13 – 30Hz; *p*_clust_ = .01).

**Figure 4.**
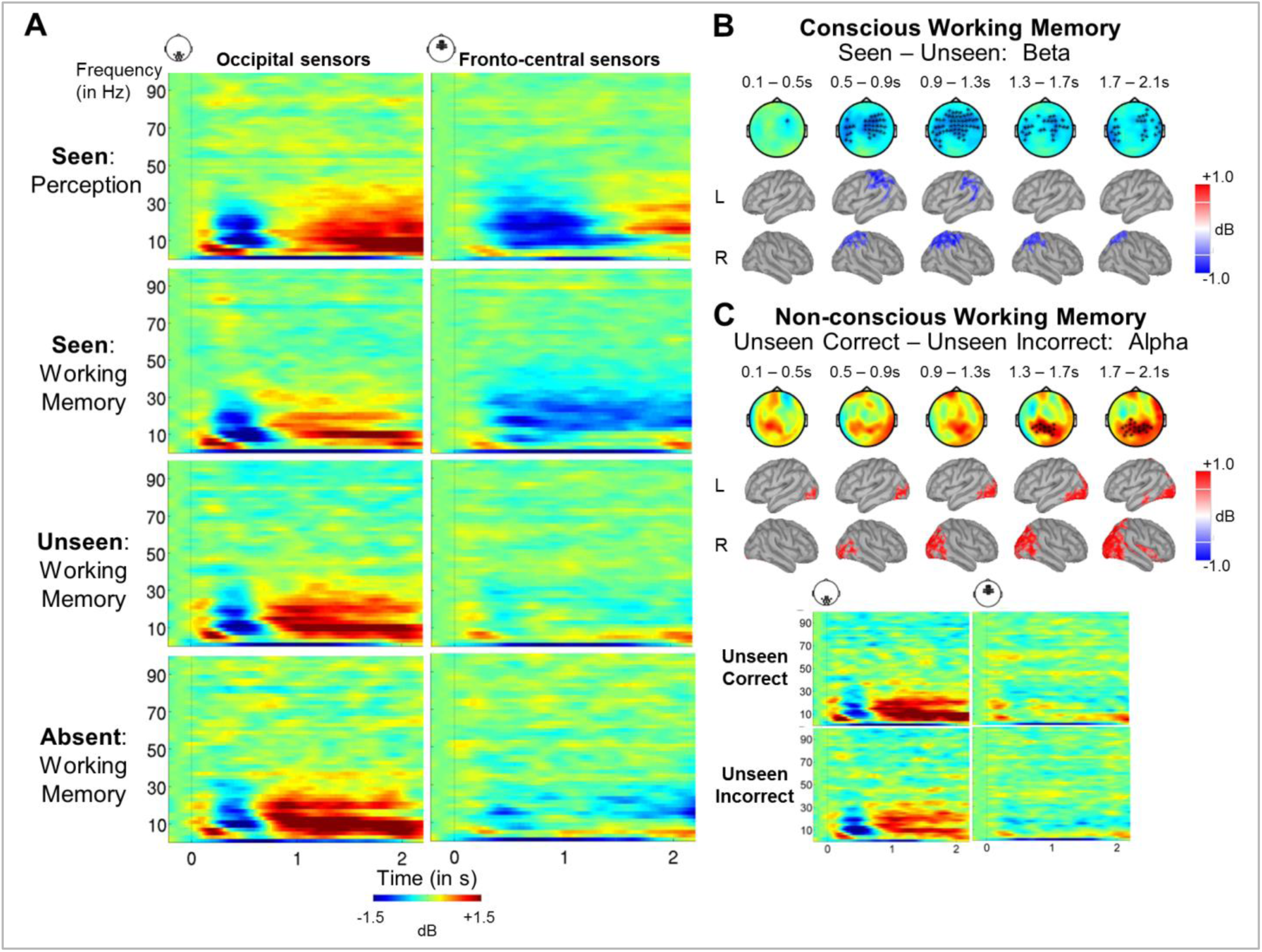
**A sustained decrease in alpha/beta power as a marker of conscious working memory**. **(A)** Average time-frequency power relative to baseline (dB) as a function of task and visibility category in a group of occipital (left) and fronto-central (right) magnetometers.**(B)** Beta band activity (13 – 30Hz; 0 – 2.5s) related to conscious working memory (seen – unseen trials) as shown in magnetometers (top) and source space (bottom; in dB relative to baseline). Black asterisks indicate sensors showing a significant difference as assessed by a Monte-Carlo permutation test.**(C)** Same as in (A) and (B) but for unseen correct and unseen incorrect trials in the alpha band (8 – 12Hz).

Most importantly, when comparing seen and unseen trials in the working memory task, we again observed a similar, but now temporally sustained, pattern of alpha/beta band desynchronization (Figure 4B). Starting at ∼300 to 500ms, seen targets evoked a power decrease in central, temporal/parietal, and frontal regions in the alpha (*p*_clust_ = .003) and beta (*p*_clus_t < .001) bands. Crucially, the beta-band desynchronization spanned the delay period and was specific to seen trials (Figure 4A), with only a couple of interspersed periods of residual desynchronization persisting in the target-absent control trials. No task- or visibility-related modulations in power spectra were found in occipital areas, and the desynchronization originated primarily from a parietal network of brain sources (Figures 4A and B). In conjunction with the afore-mentioned results, these findings confirm the major role of alpha/beta desynchronization as a correlate of conscious perception (Gaillard et al., 2009) and highlight a neural state common to conscious perception and working memory.

### A distinct neurophysiological mechanism for non-conscious working memory

Having identified a robust marker of conscious working memory, we can now evaluate the alternative hypotheses. If non-conscious working memory resulted from a small set of seen trials erroneously labeled as unseen (miscategorization hypothesis) or from the conscious maintenance of an early guess (conscious maintenance hypothesis), we would expect the same signatures as on conscious trials, including a sustained desynchronization. Conversely, the absence of such a power decrease would establish that blindsight resulted from a distinct type of non-conscious information maintenance.

There was no indication of a desynchronization when averaging over all unseen trials in the working memory task (Figure 4A). This null-effect could have reflected the mixture of trials subsumed in the unseen category, with subjects responding correctly only half of the time (RC: *M* = 49.8%, *SD* = 14.7%). Even when analyzing the unseen correct trials, however, there was no trace of any alpha/beta desynchronization (Figure 4C). Only one effect, reversed relative to conscious trials, was observed in the alpha band (*p*_clust_ = .040) in a set of posterior central sensors, corresponding to primarily occipital sources: Starting at ∼1.5s and extending until ∼1.9s, unseen correct trials exhibited a stronger *increase* in alpha power than their incorrect counterparts. These findings indicate that non-conscious working memory is a genuine phenomenon, distinct from conscious working memory.

### Contents of conscious and non-conscious working memory can be tracked transiently

We next determined where and how the specific contents of working memory were stored. Circular-linear correlations between the amplitude of the ERFs and target location (combined across all working memory trials) revealed a strong and focal association (relative to baseline) over posterior channels, starting at ∼116ms and lasting until 788ms (all *ps* < .001; Figure 5A and Table 2). Similarly, distractor position could be tracked between ∼170 and 534ms after its presentation (all *ps* < .040). The spatial position of our stimuli could thus be faithfully “decoded” in visual areas, thereby ruling out contributions from eye movements.

**Table 2.**
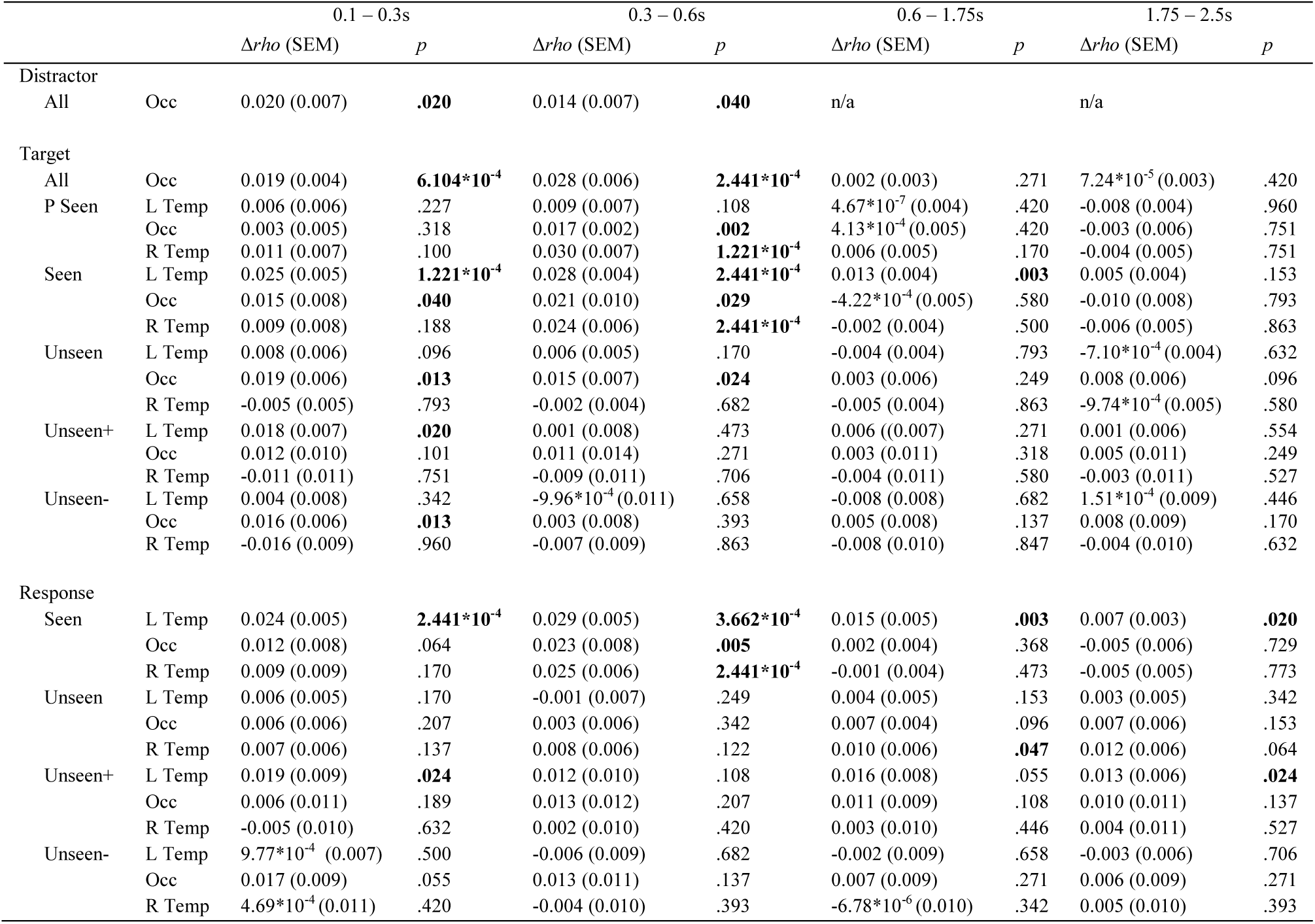
**Statistics for circular-linear correlation analyses**. Statistics for circular-linear correlation analyses between the average amplitude of the MEG signal in the gradiometers and distractor, target, and response position are listed as a function of visibility, accuracy, channel group and time window. Bold numbers indicate significant differences in correlation values relative to baseline (one-tailed Wilcoxon signed-rank test). P = perception task; Unseen+ = unseen correct trials (within +/− 2 positions of actual target location); Unseen- = unseen incorrect trials; Occ = occipital gradiometers; L Temp = left temporo-occipital gradiometers; R Temp = right temporo-occipital gradiometers; SEM = standard error of the mean (across subjects).

**Figure 5.**
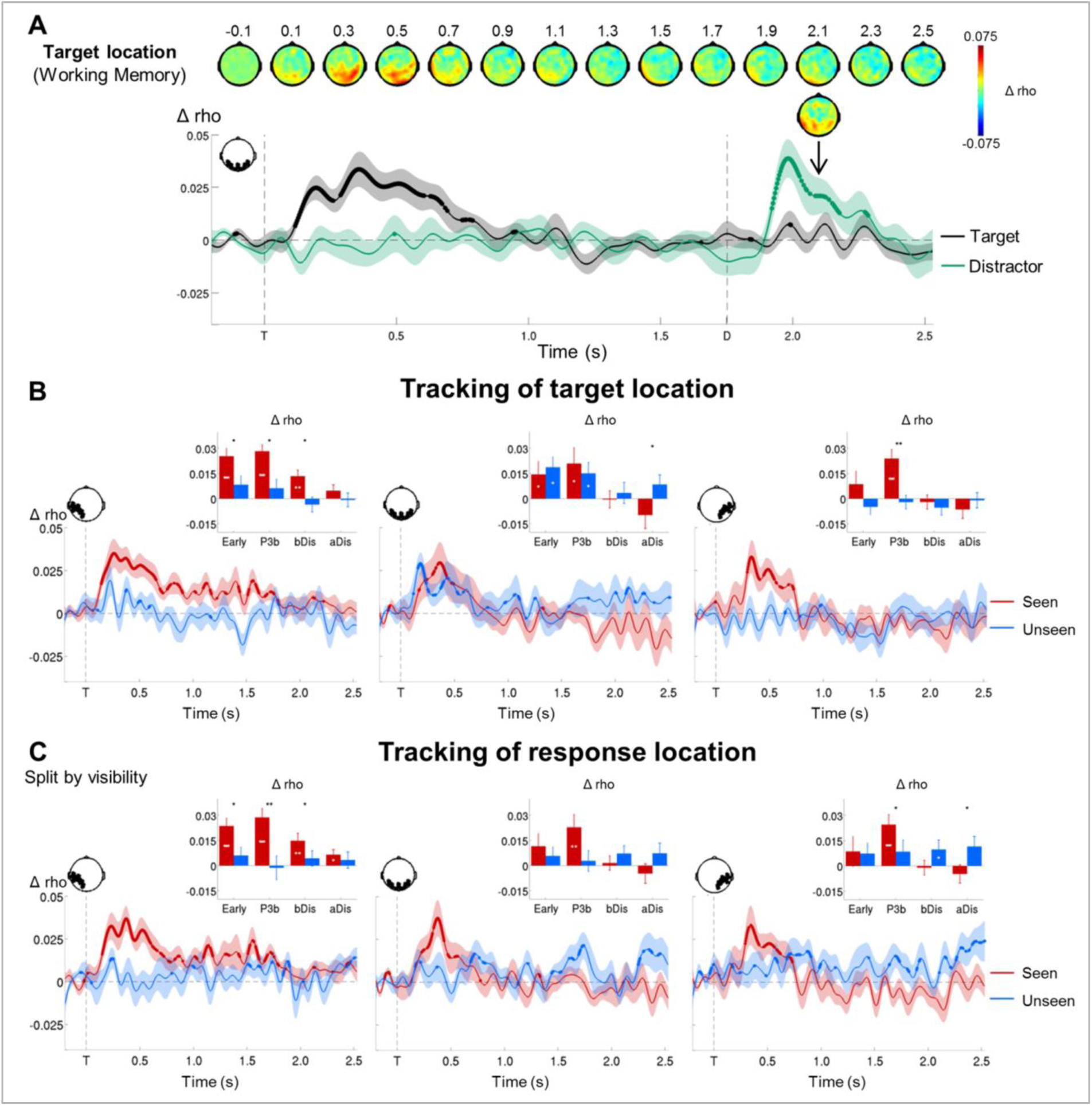
**Tracking the contents of conscious and non-conscious working memory**. **(A)** Topographies (top) and time courses (bottom; −0.2 – 2.5s) of average circular-linear correlations between the amplitude of the MEG signal (gradiometers) and target/distractor location. Shaded area demarks standard error of the mean (SEM) across subjects. Thick line represents significant increase in correlation coefficient as compared to baseline (one-tailed Wilcoxon signed-rank test across subjects, uncorrected).**(B)** Average time courses (−0.2 – 2.5s) of circular-linear correlation coefficients between amplitude of the ERFs and target location on seen trials as a function of visibility in the working memory task in a group of left temporo-occipital (left), occipital (middle), and right temporo-occipital (right) gradiometers. Shaded area demarks standard error of the mean (SEM) across subjects. Thick line represents significant increase in correlation coefficient as compared to baseline (one-tailed Wilcoxon signed-rank test across subjects,uncorrected). Insets show average correlation coefficients (relative to baseline) over four time windows: 0.1 – 0.3s (early), 0.3 – 0.6s (P3b), 0.6 – 1.75s (bDis), and 1.75 – 2.528s (aDis). White asterisks denote significant differences to baseline (one-tailed Wilcoxon signed-rank test across subjects), black asterisks significant differences between conditions (two-tailed Wilcoxon signed-rank test across subjects). For display purposes, data were lowpass-filtered at 8Hz. \**p* < .05, \*\**p* < .01, and \*\*\**p* < .001. bDis = before distractor, aDis = after distractor, T = target onset.**(C)** Same as in (B), but with response location.

In a subsequent step, we investigated how such information would be maintained in the context of conscious and non-conscious working memory (Figure 5B). Target position was weakly and transiently encoded in occipital cortex from ∼168 to 492ms on seen trials (all *ps* < .040) and, with comparable levels (all *ps* > .05), from ∼160 to 536ms on unseen trials (all *ps* < .024). In the case of seen trials, more anterior areas in left temporo-occipital cortex also coded for target location between ∼144 and 720ms and then maintained this representation in a slowly decaying, yet intermittently resurfacing manner at least up to the time of the distractor (significance was not attained in the 1.75 – 2.5s time window; all other *ps* < .003). The corresponding right-hemispheric regions encoded target position only transiently between ∼316 and 752ms (*p* < .001). Importantly, for unseen targets, neither encoding nor maintenance of location was observed in these temporo-occipital regions (all *ps* > .096). Correlation scores were always significantly higher for seen than for unseen trials (all *ps* < .05) and this absence of decodability during the maintenance period persisted, even when considering unseen correct and unseen incorrect trials separately (Supplementary Figure 1A). There was only a trace of residual decoding of target location on unseen correct trials in left temporo-occipital areas during the delay period, but this did not reach significance. Note that in the perception task, seen targets could be retrieved similarly to their counterparts in the working memory task between ∼276 and 724ms in occipital and right temporo-occipital regions (all *ps* > .244; Supplementary Figure 2). In line with previous research (Harrison and Tong, 2009; King et al., 2016), these results suggest that, in the case of conscious working memory, posterior sensory regions may encode and slightly more anterior areas maintain to-be-remembered information through a slowly decaying, intermittently reactivated, neural code (Fuentemilla et al., 2010). In contrast, no such retention mechanism appears to underlie non-conscious working memory.

### Further evidence against the conscious maintenance hypothesis

The correlation between target location and brain activity affords another way to invalidate the conscious maintenance hypothesis. If subjects quickly guessed the location of an unseen target and then held it in conscious working memory, we should be able to decode their response long before it occurs. Circular-linear correlations refuted this prediction (Figure 5C). Response position could be tracked almost identically to target location on seen (all *ps* < .020), but not on unseen trials (all *ps* > .064, with the exception of right temporo-occipital channels between 0.6 and 1.75s: *p* = .047). When we further distinguished unseen correct from unseen incorrect trials, the results remained similar, though much noisier (all *ps* > .110; Supplementary Figure 1B). There was only a hint that response position could be decoded slightly better on unseen correct than on unseen incorrect trials between 100 – 300ms and 1.75 – 2.5s in left temporo-occipital channels (*ps* < .05). Taken together, these results reject alternative explanations for the blindsight effects observed here and in previous experiments (Bergström and Eriksson, 2014; Bergström and Eriksson, 2015; Soto et al., 2011). Above-chance performance on unseen trials can neither be attributed to erroneous visibility reports, nor to the conscious maintenance of an early guess.

### Short-term synaptic change as a neurophysiological mechanism for non-conscious working memory

What mechanism might permit above-chance recall without any sustained brain activity on unseen trials? Recent modelling suggests that sustained neural firing may not be required to maintain a representation in working memory. Mongillo, Barak, and Tsodyks (2008) proposed a theoretical framework for working memory, in which information is stored in calcium-mediated short-term changes in synaptic weights, thus linking the active cells coding for the memorized item. Once these changes have occurred, the cell assembly may go dormant during the delay, while the synaptic weights are slowly decaying. At the end of the delay period, a non-specific read-out signal may then suffice to reactive the assembly. Furthermore, reactivation of the assembly may also occur spontaneously during the retention phase, similar to the rehearsal process postulated by Baddeley (2003), thus refreshing the weights and permitting the bridging of longer delays. Could this ‘activity-silent’ mechanism also constitute a plausible neural mechanism for non-conscious working memory?

To test this hypothesis, we simulated our experiments using a one-dimensional recurrent continuous attractor neural network (CANN) based on Mongillo et al. (2008). The CANN encoded the angular position of the target and was composed of neurons aligned according to their preferred stimulus value (Figure 6A). Transient short-term plasticity between the recurrent connections, with a 4s-decay constant, was implemented as described by Mongillo et al. (2008). Timing of the simulated events was comparable to the experimental paradigm: A target signal was briefly presented at a random location, followed by a mask signal to all neurons and a non-specific recall signal after a 3s-delay.

If the activity-silent mechanism constituted a plausible neurophysiological correlate of non-conscious working memory, these simulations should capture our principal findings. A stimulus presented at threshold should entail one of two different maintenance regimes: a first distinguished by near-perfect recall with spontaneous reactivations of the memorized representation throughout the retention period, and a second characterized by above-chance objective performance in the absence of delay activity.

In a noiseless model, there indeed existed a critical value of mask amplitude, *A*_critical_, which separated two distinct regimes: Just as was the case for our seen trials, when *A*_mask_ < *A*_critical_, the neural assembly coding for the target spontaneously reactivated during the delay (Figure 6C). However, when *A*_mask_ > *A*_critical_, the system evolved into a state without spontaneous activation of target-specific neurons, yet with a reactivation in response to a non-specific recall signal, mimicking our unseen trials (Figure 6D). When fixing mask amplitude near *A*_critical_ and adding noise continuously or just to the inputs, the network exhibited both types of regimes in nearly equal proportions: 50.8% of trials were characterized by spontaneous reactivations during the delay and 49.2% by an activity-silent delay period. Reminiscent of our behavioral results, sorting the trials according to the existence or absence of these reactivations and computing the histograms of recalled target position relative to true location produced two distributions of objective working memory performance: one, in which target position was nearly accurately stored (Figure 6E), and one, in which performance remained above chance despite a higher base rate of errors (Figure 6F). These simulations replicate our experimental findings and confirm that the activity-silent mechanism may underlie non-conscious working memory.

**Figure 6.**
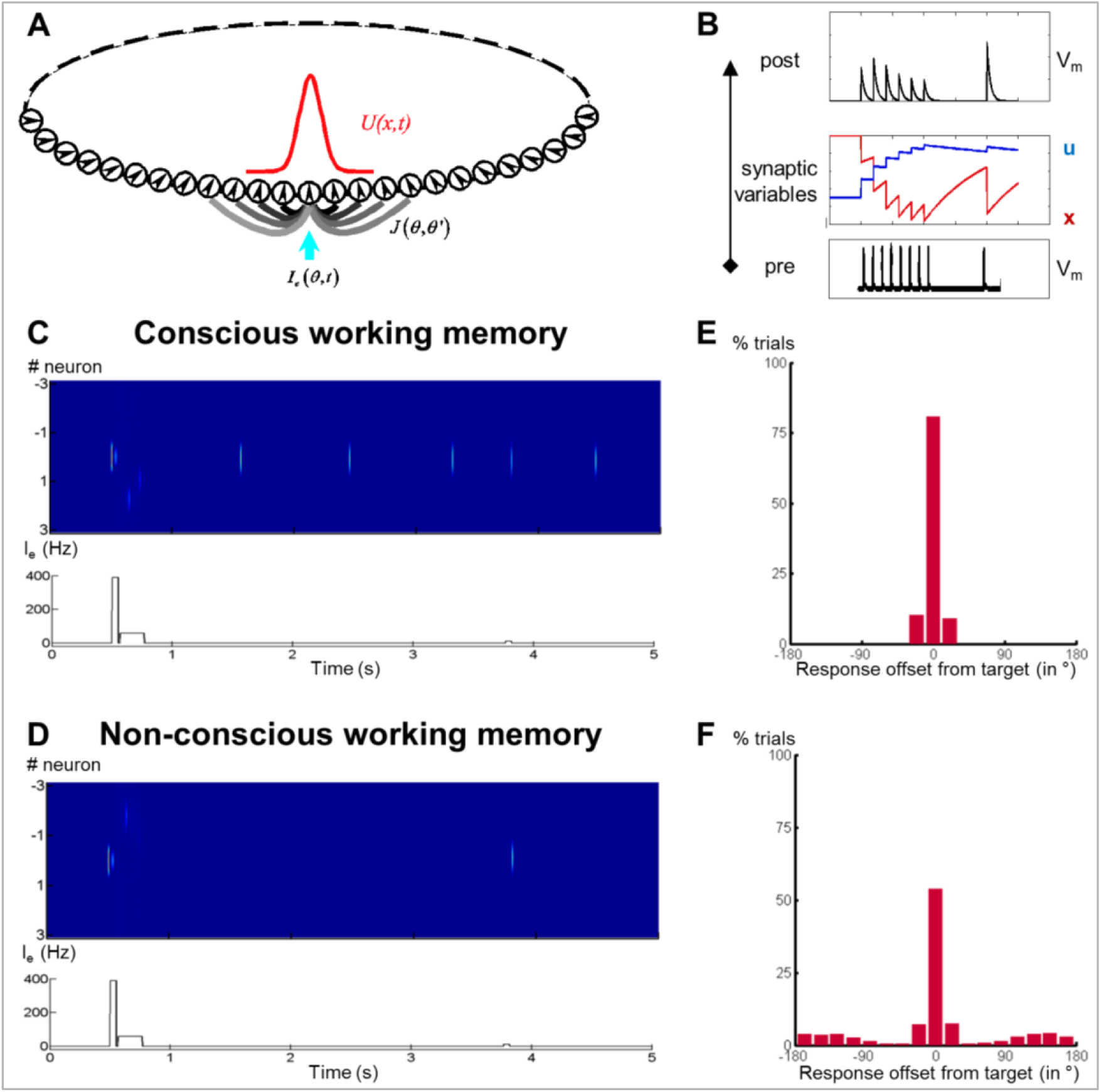
**Activity-silent neural mechanisms underlying conscious and non-conscious working memory**. **(A)** Structure of a one-dimensional continuous attractor neural network (CANN). Neuronal connections *J (θ, θ’)* are translation-invariant in the space of the neurons’ preferred stimulus values (*-π*, *π*), allowing the network to hold a continuous family of stationary states (bumps). An external input *I_e_ (θ, t)* containing the stimulus information triggers a bump state (red curve) at the corresponding location in the network.**(B)** Model of a synaptic connection with short-term potentiation. In response to a presynaptic spike train (bottom), the neurotransmitter release probability *u* increases and the fraction of available neurotransmitter *x* decreases (middle), representing synaptic facilitation and depression. Effective synaptic efficacy is proportional to *ux* (top).**(C)** Firing rate of neurons (top) and sequence of events (bottom; target and mask signal) when simulating conscious working memory with *A_mask_* = 50Hz < *A_critical_.***(D)** Same as in (C) for non-conscious working memory when *A_mask_* = 65Hz > *A_critical_.***(E, F)** Performance of the network (distribution of responses) when mask amplitude was near the critical level, *A_mask_* = 62Hz *∼A_critical_*, and noise had been added to the system. Out of 4000 trials, 2035 resulted in the conscious(E) and the remainder in the non-conscious regime (F). In both cases, performance remained above chance with the responses concentrated around the initial target location.

## Discussion

Conscious perception and working memory are thought to be intimately related, yet recent evidence challenged this assumption by suggesting the existence of non-conscious working memory (Soto et al., 2011). The present results reconcile these views. Both conscious perception and working memory shared similar mechanisms, including a beta power decrease, spanning the entire delay on working-memory trials. However, participants remained able to localize a subjectively invisible target after a 4s-delay. This non-conscious working memory could neither be explained by erroneous visibility reports nor by the conscious maintenance of an early guess. Relying on insights from a simulated neural network, we invoked an activity-silent mechanism as a potential neural correlate of non-conscious working memory. We now discuss these points in turn.

### Shared brain mechanisms underlie conscious perception and conscious working memory

Consistent with introspective reports and research on visual awareness and working memory (Baddeley, 2003; Dehaene et al., 2014), we confirmed a close relationship between conscious perception and working memory. In both tasks, classifiers trained to separate seen and unseen trials resulted in thick diagonals up to ∼1s after target onset, even when generalizing from one task to the other. Such long diagonals have repeatedly been observed in recent studies and are thought to reflect sequential processing (King and Dehaene, 2014; Marti et al., 2015; Salti et al., 2015; Stokes et al., 2015; Wolff et al., 2015). These results highlight that, irrespective of context, conscious perception involves a series of partially overlapping processing stages.

Time-frequency decompositions confirmed this conclusion. Seen trials in the perception task differed from unseen trials by a prominent decrease in alpha/beta power over fronto-central sensors, corresponding to a distributed network centered on parietal cortex. A similar desynchronization, sustained throughout the retention period, was also observed for conscious working memory. Alpha/beta band desynchronizations such as these have previously been linked with conscious perception (Gaillard et al., 2009; Wyart and Tallon-Baudry, 2009) and working memory (Lundqvist et al., 2016). Modelling suggests that the memorized item is encoded by intermittent gamma bursts, which interrupt an ongoing desynchronized beta default state (Lundqvist et al., 2011). Such a decreased rate of beta bursts, once averaged over many trials, would have resulted in the apparently sustained power decrease we observed. Increases in gamma power have also been shown in some studies on conscious perception (e.g., Gaillard et al., 2009), but we failed to detect it here (Figure 4), perhaps because our targets were brief, peripheral, and low in intensity.

Circular-linear correlations further highlighted the similarity between conscious perception and working memory. Location information could be tracked for ˜500ms on perception-only trials and for at least 1.5 seconds of the working memory retention period (Supplementary Figure 2). The mental representation formed during conscious perception was therefore either maintained or repeatedly replayed during conscious working memory.

### Long-lasting blindsight effect reflects genuine non-conscious working memory

Even when subjects indicated not having seen the target, they still identified its position much better than chance up to 4 seconds after its presentation. This long-lasting blindsight effect was replicated in two independent experiments and withstood salient visible distractors and a concurrent demand on conscious working memory. Those results corroborate previous research showing that information can be maintained non-consciously (e.g., Bergström and Eriksson, 2014; Bergström and Eriksson, 2015; Dutta et al., 2014; Soto et al., 2011). However, these prior findings could have arisen due to errors in visibility reports. If, for example, a participant had been left with a weak impression of the target (and, consequently, its location), he or she might not have had adequate internal evidence to refer to this perceptual state as seen, thus incorrectly applying the label unseen. A small number of such errors would have produced above-chance responding. Another explanation could have been the conscious maintenance of an early guess, whereby subjects would have ventured a prediction as to the correct target position immediately after its presentation and then consciously maintained this hunch.

The MEG results clearly refute these possibilities. First, whereas seen trials were characterized by a sustained desynchronization in the alpha/beta band in parietal brain areas, no such desynchronization was observed on unseen trials, even when subjects correctly identified the target location. On the contrary, the only difference between unseen correct and unseen incorrect trials emerged around the time of the distractor and was reversed in direction: Unseen correct trials were accompanied by an increase in power in the alpha band with respect to their incorrect counterpart, an effect that might relate to a successful attempt to reduce interference from the distractor (Cooper et al., 2003; Jensen and Mazaheri, 2010). Otherwise, unseen correct and incorrect trials were indistinguishable in their power spectra and similar to target-absent trials. Secondly, while target location was maintained via a slowly decaying, yet intermittently resurfacing, neural code on seen trials in posterior brain regions, there was no evidence for such maintenance-related activity on unseen trials. Circular-linear correlations between the amplitude of the MEG signal and the position of unseen targets were nevertheless initially at par with that of seen targets, but quickly dropped to baseline level. As such, the absence of delay-period activity on unseen trials does not appear to be an artifact attributable to a loss of statistical power or increase in noise. Instead, in conjunction with prior evidence (King et al., 2016; Salti et al., 2015), our findings suggest that there may be two successive mechanisms for the short-term maintenance of non-conscious stimuli: an initial, transient period of ∼1 second, during which the non-conscious representation is encoded by active firing with a slowly decaying amplitude, and a subsequent phase, during which neural activity is undetectable, yet behavior remains above chance for several seconds, thus suggesting an ‘activity-silent’ maintenance by short-term changes in synaptic weights.

### A theoretical framework for ‘activity-silent’ working memory

We presented a theoretical framework, based on Mongillo et al. (2008) and the concept of ‘activity-silent’ working memory (Stokes, 2015), that may provide a plausible explanation for the present findings. According to this model, short-term memories are maintained by slowly decaying patterns of synaptic weights. A retrieval cue presented at the end of the delay serves as a non-specific read-out signal capable of reactivating these dormant representations above chance-level. Support for this model comes from experiments in which non-specific task-irrelevant stimuli (Wolff et al., 2015) or neutral post-cues (Sprague et al., 2016) presented during a delay restore the decodability of representations. Direct physiological evidence for the postulated short-term changes in synaptic efficacies also exists (Fujisawa et al., 2008). The present non-conscious condition provides further strong support for such an activity-silent mechanism, as it was accompanied by a disappearance of delay activity (Figures 4 and 5).

In this framework, a stimulus that fails to cross the threshold for sustained activity and subjective visibility may still induce enough activity in high-level cortical circuits to trigger short-term synaptic changes. Such transient non-conscious propagation of activity has indeed been simulated in neural networks (Dehaene and Naccache, 2001) and measured experimentally in temporo-occipital, parietal, and even prefrontal cortices (Salti et al., 2015; van Gaal and Lamme, 2012). In the present work, we indeed observed some residual, transiently decodable activity over left occipito-temporal sensors on unseen correct trials (Supplementary 1). The memory of target location could therefore have arisen from posterior visual maps (Roelfsema, 2015). Note that activity-silent mechanisms need not apply solely to prefrontal cortex as originally proposed by Mongillo et al. (2008), but constitute a generic mechanism that may be replicated in different areas, possibly with increasingly longer time constants across the cortical hierarchy (Chaudhuri et al., 2014). Only some of these areas/spatial maps may be storing the information on unseen trials.

A key feature of Mongillo et al.’s (2008) model and the present simulations is that, even for above-threshold (‘seen’) stimuli, delay activity is not continuously sustained.

Occasional bouts of spontaneous reactivation instead refresh the synaptic weights and maintain the memory for an indefinite time. This account fits with Baddeley’s (2003) central hypothesis that working memory requires frequent rehearsal, and suggests that even consciously perceived items may not be “in mind” constantly. The time course of circular-linear correlations we observed on seen trials matches this description: While target location was encoded and maintained in temporo-occipital areas, target decodability was not sustained continuously, but waxed and waned throughout the delay, potentially reflecting conscious rehearsal. Fuentemilla et al. (2010) also observed that, during a delay period, decodable representations of memorized images recurred at a theta rhythm. More recently, single-trial analyses of monkey electrophysiological recordings in a working-memory task have confirmed the absence of any continuous activity and instead identified the presence of discrete gamma bursts, paired with a decrease in beta-burst probability (Lundqvist et al., 2016). Although we simulated only the simplest model of activity-silent working memory (Mongillo et al., 2008), a biologically more elaborate version (Lundqvist et al., 2011) also captures such decreases in alpha/beta power.

## Conclusion

In contrast to a widely-held belief, our findings support the existence of genuine working memory in the absence of either conscious perception or sustained activity. Following a transient encoding phase supported by active firing, non-conscious stimuli may then be maintained by ‘activity-silent’ short-term changes in synaptic weights without any detectable neural activity, allowing above-chance retrieval for several seconds. These results highlight the need to re-conceptualize our understanding of working memory, and to continuously challenge the limits of non-conscious processing.

## Methods

### Subjects

38 healthy volunteers participated in the present study (experiment 1: *N* = 17, *M*_age_ = 23.3 years, *SD*_age_ = 2.8 years, 10 men; experiment 2: *N* = 21, *M*_age_ = 24.3 years, *SD*_age_ = 3.8 years, 9 men). They gave written informed consent and received 80 or 15€ as compensation for the imaging and behavioral paradigms. Due to noisy recordings, only 13 of the 17 subjects in experiment 1 were retained for the MEG analyses.

### Experimental protocol

Participants performed variations of a spatial delayed-response task, designed to assess retention of a target location under varying levels of subjective visibility (Figure 1A). Each trial began with the presentation of a central fixation cross (500ms), displayed in white ink on an otherwise black screen. In experiment 1, a faint gray target square (RGB: 89.25 89.25 89.25) was flashed for 17ms in 1 out of 20 equally spaced, invisible positions along a circle centered on fixation (radius = 200 pixels; 8 repetitions/location). Another fixation cross (17ms) preceded the display of the mask (233ms). Mask elements were composed of four individual squares (two right above and below, and two to the left and right of the target stimulus), arranged to tightly surround the target square without overlapping it. They appeared simultaneously at all possible target locations. Mask contrast was adjusted on an individual basis in a separate calibration procedure (see below). A variable delay period with constant fixation followed the mask (experiment 1: 2.5, 3.0, 3.5, or 4.0s). On 50% of the trials in experiment 1, an unmasked distractor square, randomly placed and with the same duration as the target, was presented 1.5s into the delay period.

After the delay, 20 letters – drawn from a subset of lower-case letters of the alphabet (excluded: *e, j, n, p, t, v)* – were randomly presented in the 20 positions (2.5s). Participants were asked to identify the target location by speaking the name of the letter presented at the location. They were instructed to always provide a response, guessing if necessary. A trial ended with the presentation of the word *Vu?* (French for *seen*) in the center of the screen (2.5s), cueing participants to rate the visibility of the target on the 4-point Perceptual Awareness Scale (PAS; *1*: no experience of the target, *2*: brief glimpse, *3:* almost clear experience, *4:* clear experience; Ramsøy and Overgaard, 2004) using the index, middle, ring, or little finger of their right hand (five-button non-magnetic response box, Cambridge Research Systems Ltd., Fiber Optic Response Pad). We instructed subjects to reserve a visibility rating of *1* for those trials, for which they had absolutely no perception of the target. The target square was also replaced by a blank screen on 20% of the trials, in order to obtain an objective measure of participants’ sensitivity to the presence of the target. The inter-trial interval (ITI) lasted 1s. Subjects completed a total of 200 trials of this working memory task, divided into four separate experimental blocks. They also undertook two blocks of 100 trials of a perception-only control paradigm, identical to the working memory task in all respects except that the delay period and target localization screen were omitted, such that the presentation of the mask immediately preceded subjects’ visibility ratings. Task order (perception vs. working memory) was counterbalanced across participants.

Experiment 2 was designed to investigate the impact of a conscious working memory load on non-conscious working memory. Apart from the following exceptions, it was identical to experiment 1: A screen with either 1 (low load) or 5 (high load) centrally presented digits (1.5s) – randomly drawn (without replacement) from the numbers 1 through 9 – as well as a 1s-fixation period were shown prior to the presentation of the target square. Following either a 0s-or a 4s-delay period, subjects first identified the target location by typing their responses on a standard AZERTY keyboard (4s). The French word for *numbers* (*Numéros?*) then probed participants to recall the sequence of digits in the correct order. Responses were again logged on the keyboard during a period of 4.5s. Subjects last rated target visibility as in experiment 1 (3s). The ITI varied between 1 and 2s. Participants completed two experimental blocks of 100 trials each.

### Calibration task

Prior to the experimental tasks, each participant’s perceptual threshold was estimated in order to ensure roughly equal proportions of seen and unseen trials. Subjects completed 150 (experiment 1: 3 blocks) or 125 (experiment 2: 5 blocks) trials of a modified version of the working memory task (no distractor, delay duration: 2s in experiment 1 and 0s in experiment 2), during which mask contrast was either increased (following a visibility rating of *2*, *3*, or *4*) or decreased (following a visibility rating of *1*) on each target-present trial according to a double-staircase procedure. Individual perceptual thresholds to be used in the main tasks were derived by averaging the mask contrasts from the last four switches from seen to unseen (or vice versa) in each staircase.

### Behavioral analyses

We analyzed our behavioral data in Matlab R2014a (MathWorks Inc., Natick, MA) and SPSS Statistics Version 20.0 (IBM, Armonk, NY). Only meaningful trials without missing responses were included in any analysis. Distributions of localization responses were computed for visibility categories with at least five trials per subject. Objective working memory performance was quantified via two complementary measures. The *rate of correct responding* (CR) was defined as the proportion of trials within two positions (i.e.,+/− 36°) of the actual target location and served as an index of the amount of information that could be retained. Because 5 out of 20 locations were counted as correct, chance on this measure was 25%. The *precision* of working memory was estimated as the dispersion (standard deviation) of spatial responses. In particular, we modeled the observed distribution of responses *D(n)* as a mixture of a uniform distribution (random guessing) and an unknown probability distribution *d* (“true working memory”):

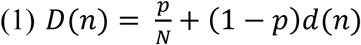

where *p* refers to the probability that a given trial is responded to using random guessing; *N* to the number of target locations (*N* = 20); and *n* is the deviation from the true target location. We assumed that *d*(*n*) = 0 for deviations beyond a fixed limit *a* (with *a* = 2). This hypothesis allowed us to estimate *p* from the mean of that part of the distribution *D* for which one may safely assume no contribution of working memory:

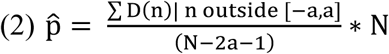

where the model is designed in such a way as to ensure that 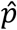 = 1 if *D* is a uniform distribution (i.e., 100% of random guessing) and 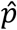 = 0 if *D* vanishes outside the region of correct responding (i.e., 0% of random guessing). There needs to be at least chance performance inside the region of correct responding, so

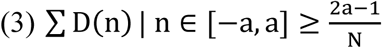

which ensures 0 ≤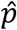 ≤ 1. This is the reason why, when computing precision, we included only subjects whose CR for unseen trials, collapsed across all experimental conditions, significantly exceeded chance performance (i.e., 25%) in a *χ*^2^-test (*p* < .05). An estimate of *d*, 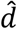, can then be derived in two steps from Equation 1 as

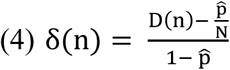

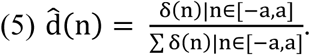

We note that the distribution δ has residual, yet negligible, positive and negative mass (due to noise) outside the region of correct responding. In order to obtain 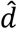, we therefore restricted the distribution δ to [*−a, a*], set all negative values to 0, and renormalized its mass to 1. The precision of the representation of the target location in working memory was then defined as the standard deviation of that distribution.

### MEG recordings and preprocessing

In experiment 1, we recorded MEG with a 306-channel (102 sensor triplets: 1 magnetometer and 2 orthogonal planar gradiometers), whole-head setup by ElektaNeuromag^®^ (Helsinki, Finland) at 1000Hz with a hardware bandpass filter between 0.1 and 330 Hz. Eye movements as well as heart rate were monitored with vertical and horizontal EOG and ECG channels. Prior to installation of the subject in the MEG chamber, we digitized three head landmarks (nasion and pre-auricular points), four head position indicator (HPI) coils placed over frontal and mastoïdian skull areas, and 60 additional locations outlining the participant’s head with a 3-dimensional Fastrak system (Polhemus, USA). Head position was measured at the beginning of each run.

Our preprocessing pipeline followed Marti et al. (2015). Using MaxFilter Software (ElektaNeuromag®, Helsinki, Finland), raw MEG signals were first cleaned of head movements, bad channels, and magnetic interference originating from outside the MEG helmet (Taulu et al., 2004), and then downsampled to 250Hz. We conducted all further preprocessing steps with the Fieldtrip toolbox (http://www.fieldtriptoolbox.org/; Oostenveld et al., 2011) run in a Matlab R2014a (MathWorks Inc., Natick, MA) environment. Initially, MEG data were epoched between −0.5 and +2.5s with respect to target onset. Trials contaminated by muscle or other movement artifacts were then identified and rejected in a semi-automated procedure, for which the variance of the MEG signals across sensors served as an index of contamination. To remove any residual eye-movement and cardiac artifacts, we performed independent component analysis separately for each channel type, visually inspected the topographies and time courses of the first 30 components, and subtracted any contaminated component from the MEG data. Epochs retained for all investigations based on evoked responses were then low-pass filtered at 30Hz. Following King et al. (2016), to track the neural representations of target, response, and distractor location, these filtered epochs were transformed into circular-linear correlation coefficients by combining the two linear correlation coefficients between the MEG signal and the sine and cosine of the angle defining the target location. A sliding, frequency-independent Hann taper (window size: 500ms, step size: 20ms) was convolved with the unfiltered epochs in order to extract an estimate of power between 1 and 99Hz (in 2Hz steps) to identify the neural correlates of conscious and non-conscious perception and working memory in the frequency domain. Prior to univariate or multivariate statistical analysis, data (ERFs, circular-linear correlation coefficients, time-frequency power estimates) were baseline corrected using a period between −200 and −50ms.

### Sources

Individual anatomical magnetic resonance images (MRI), obtained with a 3D T1-weighted spoiled gradient recalled pulse sequence (voxel size: 1 * 1 * 1.1mm; repetition time [TR]: 2,300ms; echo time [TE]: 2.98ms; field of view [FOV]: 256 * 240 * 176mm; 160 slices) in a 3T Tim Trio Siemens scanner, were first segmented into gray/white matter as well as subcortical structures with FreeSurfer (http://surfer.nmr.mgh.harvard.edu/). We then reconstructed the cortical, scalp, and head surfaces in Brainstorm (http://neuroimage.usc.edu/brainstorm; Tadel et al., 2011) and co-registered these anatomical images with the MEG signals, using the HPI coils and the digitized head shape as a reference. Current density distributions on the cortical surface were subsequently estimated separately for each condition and subject. Specifically, we employed an analytical model with overlapping spheres to compute the leadfield matrix and modeled neuronal current sources with an unconstrained (dipole orientation loosening factor: 0.2) weighted minimum-norm current estimate (wMNE; depth-weighting factor: 0.5) and a noise covariance obtained from the baseline period of all trials. Average time-frequency power in the alpha (8 – 12Hz) and beta (13 – 30 Hz) bands was then estimated with complex Morlet wavelets using the Brainstorm default parameters, the resulting transformations projected onto the ICBM 152 anatomical template (Fonov et al., 2011, 2009), and the contrasts between the conditions of interest computed. Group averages for spatial clusters of at least 150 vertices are shown in dB relative to baseline and were thresholded at 60% of the maximum amplitude (cortex smoothed at 60%).

### Multivariate pattern analyses

We employed the Scikit-Learn package (Pedregosa et al., 2011) as implemented in MNE 0.13 (Gramfort, 2013; Gramfort et al., 2014) in order to conduct our multivariate pattern analyses (MVPA). Following Marti et al. (2015), we trained a linear support vector machine (Chang and Lin, 2011) at each time sample within each participant to isolate the topographical patterns best differentiating seen and unseen trials separately for each task. Using a 5-fold, stratified cross-validation procedure, the MEG data were first randomly split into five sets of trials with the same proportion of samples for each class. 50% of the most informative features (i.e., channels) for each fold were then selected by means of a simple, univariate analysis of variance (Charles et al., 2014; Haynes and Rees, 2006) to reduce the dimensionality of the data, the remaining channel-time features *z*-score normalized, and a weighting procedure applied in order to counteract the effects of any class imbalances. The classifier was then trained on four of these sets and tested on the left-out trials in order to identify the hyperplane (i.e., topography) best suited to separate the two classes while avoiding overfitting. This sequence of events (univariate feature selection, normalization, training and testing) was repeated five times, ensuring that each trial would be included in the test set once.

Within the same cross-validation loop, we also evaluated the ability of each classifier to discriminate visibility ratings at all other time samples (i.e., generalization across time). This kind of MVPA results in a temporal generalization matrix, in which each entry represents the decoding performance of each classifier trained at time point *t* and tested at time point *t’*, and in which the diagonal corresponds to classifiers trained and tested on the same time points (King and Dehaene, 2014). Importantly, when assessing the similarity of the neural processes involved in conscious perception and working memory and thus when interrogating the capacity of our classifiers to generalize across tasks (i.e., from the perception to the working memory task and vice versa), we modified the aforementioned cross-validation procedure to capitalize on the independence of our training and testing data (see http://martinos.org/mne/dev/auto_examples/decoding/plot_decoding_time_generalization_conditions.html#example-decoding-plot-decoding-time-generalization-conditions-py as an example). As such, classifiers from each training set were directly applied to the entire testing set and the respective predictions averaged.

Classifiers generated a continuous output in the form of the distance between the respective sample and the separating hyperplane for each test trial. In order to be able to compare classification performance across subjects, we then applied a receiver operating characteristic analysis across trials within each participant and summarized overall effect sizes with the non-parametric area under the curve (AUC). Unlike average decoding accuracy, the AUC serves as an unbiased measure of decoding performance as it represents the true-positive rate (e.g., a trial was correctly categorized as seen) as a function of the false-positive rate (e.g., a trial was incorrectly categorized as seen). Chance performance, corresponding to equal proportions of true and false positives, therefore leads to an AUC of 0.5. Any value greater than this critical level implies better-than-chance performance, with an AUC of 1 indicating a perfect prediction for any given class.

### Statistical analyses

We performed statistical analyses across subjects. For the ERF and time-frequency data, cluster-based, non-parametric *t*-tests with Monte Carlo permutations were used to identify significant differences between experimental conditions (Maris and Oostenveld, 2007). Further planned comparisons of ERF time courses (seen vs. unseen) in a-priori defined spatio-temporal regions of interest (i.e., P3b time window: 300 – 600ms) were conducted with non-parametric signed-rank tests (*p*_uncorrected_ < .05). A correction for multiple comparisons was then applied with a false discovery rate (*p*_FDR_ < .05).

Non-parametric signed-rank tests (*p*_uncorrected_ < .05) were also employed to evaluate decoding performance and the strength of circular-linear correlations. Specifically, we assessed whether classifiers could predict the trials’ classes better than chance (AUC > 0.5) and whether circular-linear correlation coefficients deviated from baseline values (Δ rho > 0). We report temporal averages over four a-priori time bins, corresponding to an early perceptual period (0.1 – 0.3s), the P3b time window (0.3 – 0.6s), and the delay before (0.6 – 1.75s) and after (1.75 – 2.53s) the distractor. To capitalize on the increased spatial selectivity of gradiometers, averaged time courses of these two channels are shown for circular-linear correlations.

### Simulations

A one-dimensional, recurrent continuous attractor neural network (CANN) model (Mongillo et al., 2008) was adapted in order to simulate the experimental findings (Figure 6A). Individual neurons were aligned according to their preferred stimulus value, enabling the network to encode angular position of a target stimulus (range: -π to π; periodic boundary condition). The dynamics of this system were determined by the synaptic currents of each neuron given by

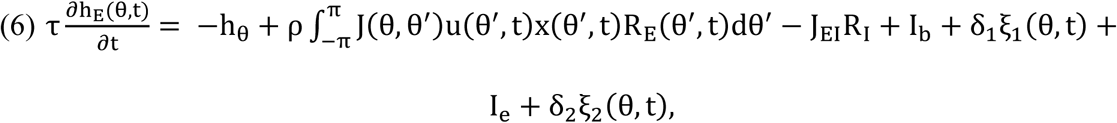

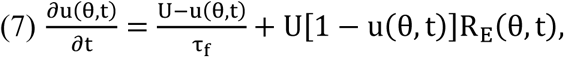

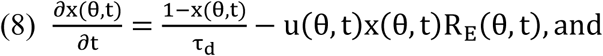

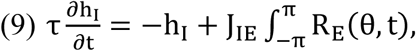

where *τ* describes the time constant of firing rate dynamics (in the order of milliseconds); *ρ* refers to neuronal density; *h_E_* (*θ,t*) and *R_E_* (*θ,t*) capture the synaptic current to and firing rate of neurons with preference θ at time *t* respectively; and *R*(*h*) = *α* ln(1 + exp(*h*/*α*)) is the neural gain chosen in the form of a smoothed threshold-linear function. *J_IE_* and *J_EI_* represent the connection strength between excitatory and inhibitory neurons. All excitatory neurons received a constant background input, *I_e_*, reflecting the arousal signal when the neural system was engaged in a working memory task. *δ_1_ξ_1_* is background noise; *I_e_*, any external stimulus (e.g., target, mask, and recall signal); and *δ_1_ξ_1_* (*t*) the noise related to those external stimuli. *u* (*θ*, *t*) and *x* (*θ*, *t*) denote the short term synaptic facilitation (STF) and depression (STD) effects at time *t* of neurons with preference *θ*, respectively. The short term plasticity dynamics are characterized by the following parameters: *J_1_* (absolute efficacy), *U* (increment of the release probability when a spike arrives), *τf* and *τ_d_* (facilitation and depression time constants). The STF value *u* (*θ*, *t*) is facilitated whenever a spike arrives, and decays to the baseline *U* within the time *τf*. The neurotransmitter value *x* (*θ*, *t*) is utilized by each spike in proportion to *u* (*θ*, *t*) and then recovers to its baseline, 1, within the time τ_*d*_.

*J* (*θ, θ*’) is the interaction strength from neurons at *θ* to neurons at *θ ’* and is chosen to be

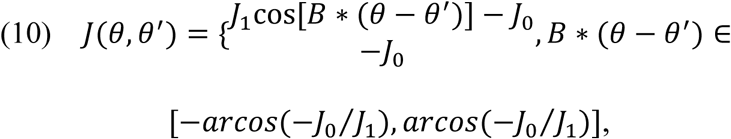

where *J_0_, J_1_*, and *B* are constants which determine the connection strength between the neurons. Note that *J* (*θ*, *θ’)* is a function of *θ* – *θ’*, i.e., the neuronal interactions are translation-invariant in the space of neural preferred stimuli. The other parameters of the system were as follows: τ = 0.008s, τ _*f*_ = 4s, τ _*d*_ = 0.3s, *J1* = 12, *J_0_* = 1, *J_EI_* = 1.9, *J_IE_* = 1.8, *I_b_* =- 0.1 Hz, *δ*_1_ = 0.3, *δ*_2_ = 9, *N* = 100, *α* = 1.5, *B* = 2.2.

During our simulations, we first presented a target signal with an amplitude of *A*_target_ = 390Hz at a random location (50ms), waited for 17ms, and then applied a mask signal to all the neurons in the system (200ms). The amplitude of the mask signal was initially varied in order to determine a critical value which would produce two distinct maintenance patterns, but was then fixed at a threshold of *A*_mask_ = 62Hz. At the end of a 3s-delay period, a non-specific recall signal was given for 50ms with *A*_recall_ = 10Hz. Remembered target position was calculated as the population vector angle during this time period.

## Acknowledgements

This work was supported by CEA, INSERM, Collège de France, a senior ERC grant – NeuroConsc – to S.D., and the Fondation Roger de Spoelberch. D.T. was funded by the ENP and the Fondation Schneider Electric. We gratefully acknowledge Henrik Ueberschär, Leila Azizi, and Virginie Van Wassenhove for their invaluable daily support and stimulating discussion.

## Supplementary Figures

**Supplementary Figure 1.**
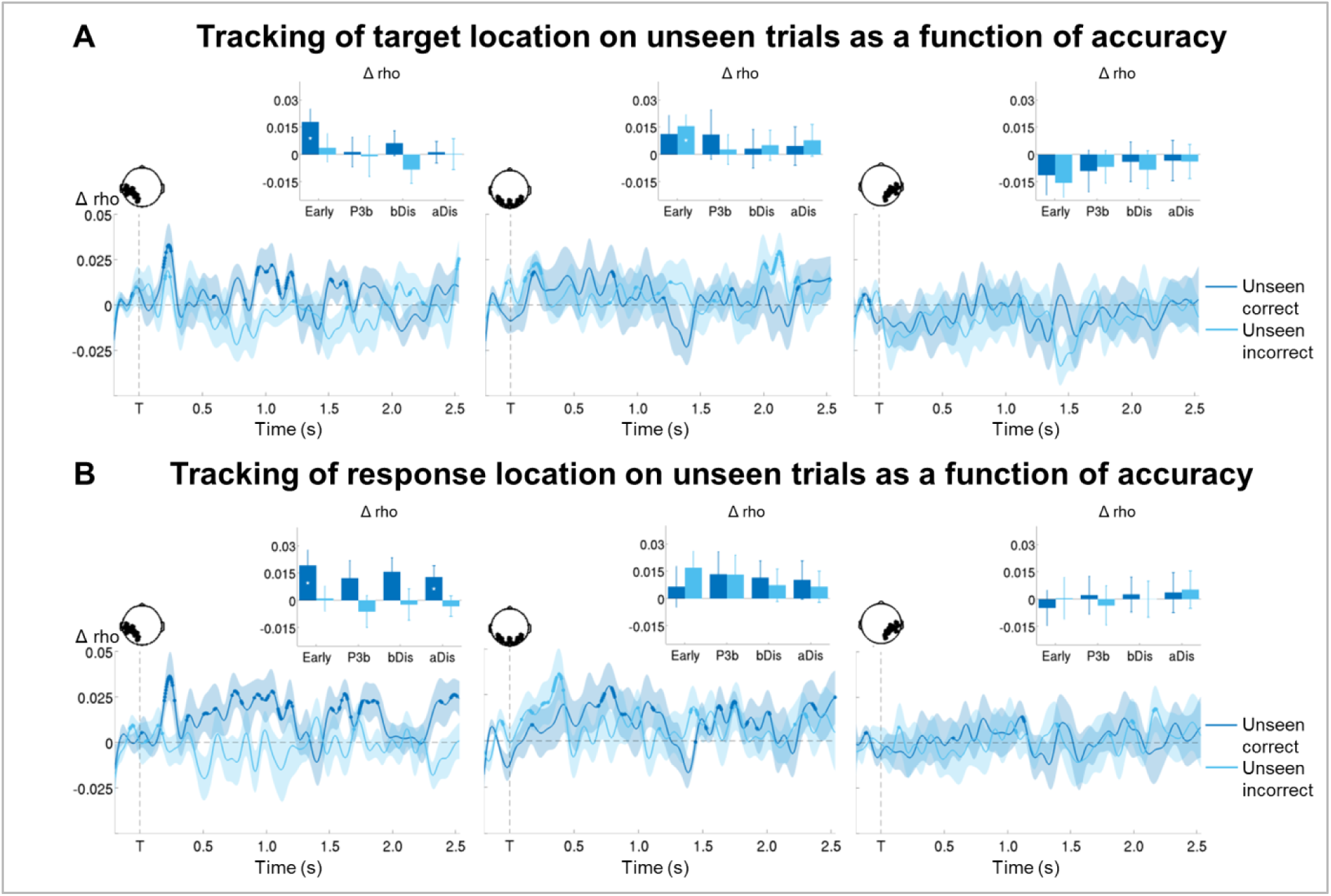
**Tracking the contents of non-conscious working memory**. **(A)** Average time courses of circular-linear correlation coefficients with target location in the working memory task (−0.2 – 2.5s) on unseen trials as a function of accuracy (correct = within +/− 2 positions of actual target location) in a group of left temporo-occipital (left), occipital (middle), and right temporo-occipital (right) gradiometers. Thick lines indicate significant increases in correlation coefficients as compared to baseline (one-tailed Wilcoxon signed-rank test across participants, uncorrected). Shaded area illustrates standard error of the mean (SEM) across subjects. Insets show average correlation coefficients (relative to baseline) over four time windows: 0.1 – 0.3s (early), 0.3 – 0.6s (P3b), 0.6 – 1.75s (bDis), and 1.75 – 2.528s (aDis). White asterisks denote significant differences to baseline (one-tailed Wilcoxon signed-rank test across participants), black asterisks significant differences between conditions (two-tailed Wilcoxon signed-rank test across subjects). For display purposes, data were lowpass-filtered at 8Hz. \**p* < .05, \*\**p* < .01, and \*\*\**p* < .001. T = target onset, bDis = before distractor, aDis = after distractor.**(B)** Same as in (A), but with response location.

**Supplementary Figure 2.**
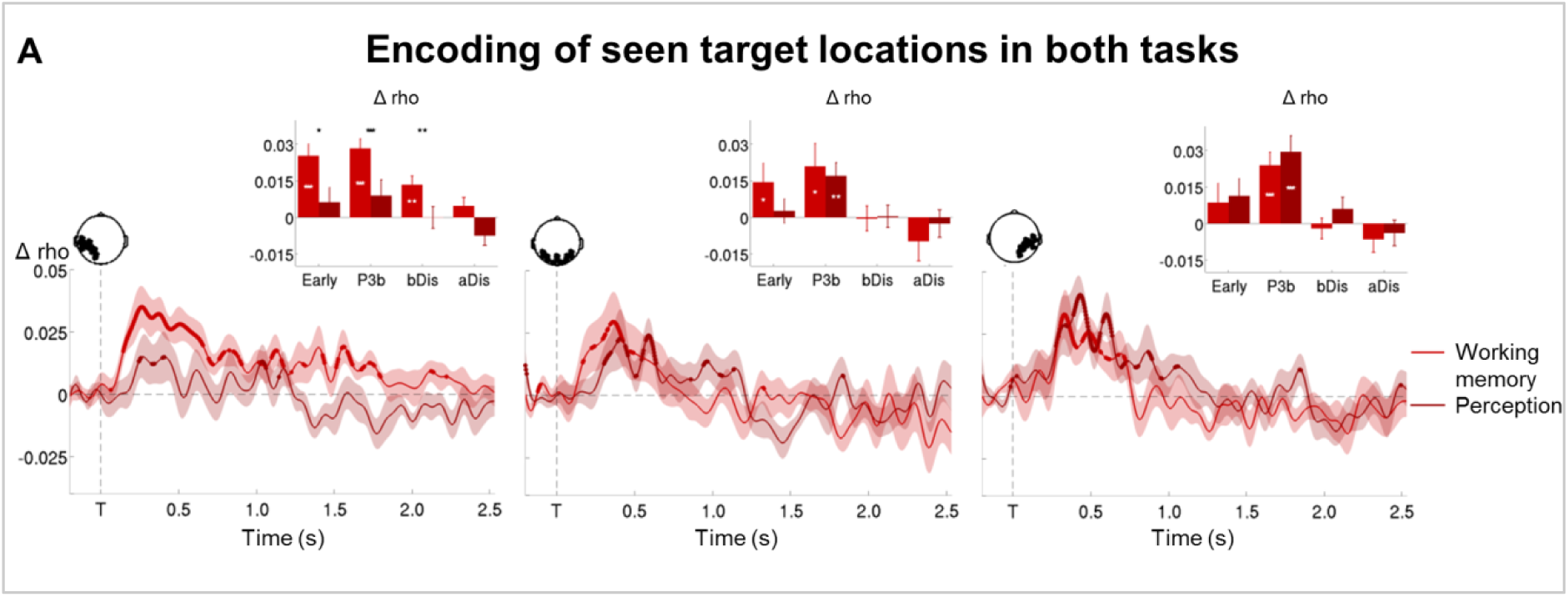
**Representation of seen target locations during conscious perception and working memory**. **(A)** Average time courses of circular-linear correlation coefficients between amplitude of the ERFs and target location on seen trials as a function of task (perception and working memory) in a group of left temporo-occipital (left), occipital (middle), and right temporo-occipital (right) gradiometers. Shaded area demarks standard error of the mean (SEM) across subjects. Thick line represents significant increase in correlation coefficient as compared to baseline (one-tailed Wilcoxon signed-rank test across subjects, uncorrected). Insets show average correlation coefficients (relative to baseline) over four time windows: 0.1 – 0.3s (early), 0.3 – 0.6s (P3b), 0.6 – 1.75s (bDis), and 1.75 – 2.528s (aDis). White asterisks denote significant differences to baseline (one-tailed Wilcoxon signed-rank test across subjects), black asterisks significant differences between conditions (two-tailed Wilcoxon signed-rank test across subjects). For display purposes, data were lowpass-filtered at 8Hz. *\*p* < .05, *\*\*p* < .01, and *\*\*\*p* < .001. T = target onset, bDis = before distractor, aDis = after distractor.

